# Physcomitrella SUN2 mediates MTOC association to the nuclear envelope and facilitates chromosome alignment during spindle assembly

**DOI:** 10.1101/2023.03.29.534829

**Authors:** Mari W. Yoshida, Noiri Oguri, Gohta Goshima

## Abstract

Plant cells lack centrosomes and instead utilise acentrosomal microtubule organising centres (MTOCs) to rapidly increase the number of microtubules at the onset of spindle assembly. Although several proteins required for MTOC formation have been identified, how the MTOC is positioned at the right place is not known. Here, we show that the inner nuclear membrane protein SUN2 is required for MTOC association with the nuclear envelope (NE) during mitotic prophase in the moss *Physcomitrium patens*. In actively dividing protonemal cells, microtubules accumulate around the NE during prophase. In particular, regional MTOC is formed at the apical surface of the nucleus. However, microtubule accumulation around the NE was impaired and apical MTOCs were mislocalised in *sun2* knockout (KO) cells. In addition, chromosome distribution in the nucleus was skewed, suggesting that SUN2 mediates the linking of microtubules with chromosomes. Upon nuclear envelope breakdown (NEBD), the mitotic spindle was assembled with mislocalised MTOC, which were a source of microtubules in *sun2* KO plants. However, completion of chromosome alignment in the spindle was delayed; in severe cases, the chromosome was transiently detached from the spindle body. SUN2 tended to localise to the apical surface of the nucleus during prophase in a microtubule-dependent manner. Based on these results, we propose that SUN2 facilitates the attachment of microtubules to chromosomes during spindle assembly by linking them prior to NEBD. Furthermore, this study suggests that trans-NE microtubule-chromosome linking, a well-known function of SUN in animals and yeast, is conserved in plants.

## Introduction

The linking of the nucleus to the cytoskeleton is a common feature of eukaryotic cells. The nuclear-associated cytoskeleton determines nuclear position, which is involved in cell physiology and fate, and applies force to the nucleus, which affects gene expression (Almonacid et al., 2019; Gundersen and Worman, 2013). Linkage between the nucleus and the microtubule organising centre (MTOC) has been reported in animals and fungi; centrosomes in certain animal cell types and the spindle pole body (SPB) in yeast are associated with the nuclear envelope (NE) (Mejat and Misteli, 2010). The key conserved factors that link the nucleus to the cytoskeleton in animals and yeasts is the linkers of the nucleoskeleton to the cytoskeleton (LINC) complex, which comprises the inner nuclear membrane protein SUN and the outer nuclear membrane protein KASH (Jahed et al., 2021; Meier, 2016; Mejat and Misteli, 2010). SUN has three recognisable domains: a transmembrane region, multimerisation domain, and C-terminal SUN domain. The SUN domain binds to the C-terminus of KASH in the nuclear intermembrane region, whereas the N-terminus of SUN interacts with chromosomes. The N-terminus of the KASH protein interacts with microtubules and/or actin filaments in the cytoplasm either directly or via motor proteins (myosin, kinesin, and dynein). Mutations in SUN in animals cause a variety of cellular defects, such as nuclear deformation and mispositioning, and sometimes cause diseases in humans (Meinke et al., 2014; Mejat and Misteli, 2010). In yeast, mutations in SUN lead to lethality owing to the failure of SPB separation and bipolar spindle formation during mitosis (Hagan and Yanagida, 1995; Jaspersen et al., 2006). The role of SUN in telomere anchorage during meiosis is also conserved in yeast and animals (Mejat and Misteli, 2010).

In plants, the SUN family is classified as Cter-SUN or mid-SUN depending on whether the SUN domain is located near the C-terminus, similar to animal and fungal SUNs, or in the middle of the protein (Graumann et al., 2014). Similar to animal/yeast orthologues, Cter-SUN is an inner nuclear membrane protein required for nuclear morphology, movement, and telomere anchorage during meiosis (Meier et al., 2017). In *AtSUN1* mutant, the shape of the nucleus is circular (Oda and Fukuda, 2011; Zhou et al., 2012). In contrast, the plant-unique mid-SUN is localised not only in the NE, but also in the ER (Graumann et al., 2014). Arabidopsis mid-SUNs (SUN3, 4, 5) are redundantly essential for early seed development and are involved in nuclear morphology (Graumann et al., 2014). Regarding the link to the cytoskeleton, nuclear migration in Arabidopsis is driven by actin and myosin XI-i, which bind to the WIT-WIP complex, i.e., the functional homologue of KASH, which interacts with SUN (Tamura et al., 2013). However, whether the SUN-KASH bridge is linked to the microtubule cytoskeleton in plants remains unclear.

The moss *Physcomitrium patens* is a model system suitable for studying cytoskeletal and nuclear dynamics, owing to its amenability to high-resolution live microscopy and gene editing techniques. Recent studies have revealed that microtubules and two kinesin family proteins drive nuclear migration (KCH for retrograde migration and ARK for anterograde migration) (Yamada and Goshima, 2018; Yoshida et al., 2023). However, the mechanisms by which KCH and ARK recognise the nucleus remain unclear. We initiated the current study to test the hypothesis that the SUN-KASH-kinesin axis is responsible for nuclear motility. First, we attempted to conduct a loss-of-function study of *SUN* genes, which are more easily identifiable via sequence homology searches than *KASH/WIP/WIT* genes. In addition to nuclear motility, we observed defects in MTOC and chromosomal positioning in prophase following the deletion of *SUN2*, one of two Cter-SUN. Moreover, the *sun2* knockout (KO) line showed delayed chromosome congression during prometaphase. Thus, this study showed that SUN is involved in MTOC positioning and attachment to the NE, revealing the functional conservation of SUN in the three major kingdoms. Furthermore, the data suggest that microtubule attachment to the chromosome during spindle assembly is facilitated by trans-NE linkage, which is mediated by SUN2 at a prior stage.

## Results

### Nuclear migration in subapical cells is suppressed in the absence of SUN2

*P. patens* possesses two Cter-SUNs (SUN1 and SUN2) and two mid-SUNs (SUN3 and SUN4) (Fig. 1A). To investigate the function of Cter-SUN in *P. patens*, we aimed to delete nearly the entire open reading frame (ORF) of *SUN1* and *SUN2* using CRISPR/Cas9 in a line expressing GFP-tubulin and histoneH2B-mCherry. We successfully obtained knockout (KO) lines for *SUN2* (Fig. S1A, B). The *sun2* KO line grew indistinguishably from the parental line on culture plates (Fig. 1B, C).

**Figure 1.**
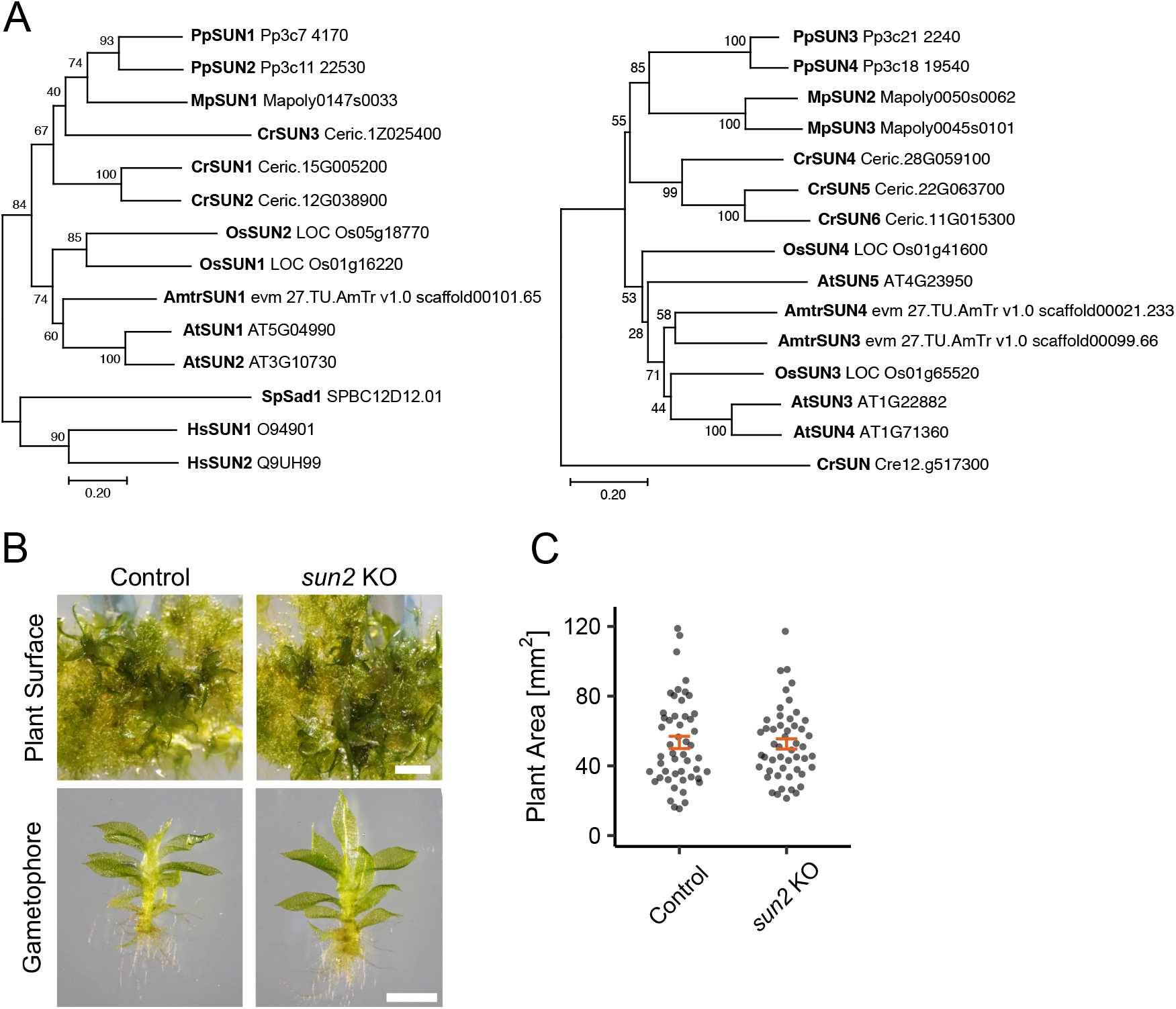
Normal development of *Physcomitrium patens sun2* knockout line. (A)Phylogenetic analysis of Cter-*SUN* genes (left) and mid-*SUN* genes (right): moss *Physcomitrium patens* (Pp), Brassica *Arabidopsis thaliana* (At), rice *Oryza sativa* (Os), green alga *Chlamydomonas reinhardtii* (Cr), *Amborella trichopoda* (Amtr), liverwort *Marchantia polymorpha* (Mp), yeast *Schizosaccharomyces pombe* (Sp), and *Homo sapiens* (Hs). Amino acid sequences were collected from the database (accession numbers are indicated on the right), aligned with MAFFT, and gaps were deleted. The phylogenetic tree was constructed using the neighbour-joining method and MEGAX software, and its reliability was assessed using 1,000 bootstrapping trials. The bar indicates 0.2 amino acid substitutions per site. (B) (Top) Culture plate containing 20-day-old moss that grew from a piece of protonemata. (Bottom) Isolated gametophores and rhizoids. Control; GFP-α-tubulin/Histone-mCherry line. Bars, 1 mm. (C) Plant area comparison. The moss lines used in this analysis are the same as those used in (B). The mean area (mm2) was 53.4 ± 3.53 (control, ±SEM, *n* = 50) and 52.6 ± 2.93 (*sun2 KO*, ±SEM, *n* = 50). P = 0.9259 based on the two-sided Mann-Whitney U test.

To identify the phenotypes of the *sun2* KO line at the cellular level, we conducted long-term time-lapse imaging of the microtubules and chromosomes using low-resolution microscopy (Fig. 2A; Movie 1). Changes in nuclear morphology were not convincingly detected by this microscopy. In contrast, abnormal nuclear movement was observed. In the wild-type protonemata, the daughter nuclei moved in the apical direction in the apical daughter cell and in the basal direction in the subapical daughter cell after apical cell division (Fig. 2A, B). In the *sun2* KO line, nuclear movement of apical cells was comparable to that of control cells. However, the rate of basal motility significantly decreased in the subapical cells from 145 min after anaphase onset (Fig. 2B, C). In addition, abnormal apical movement was occasionally observed (Fig. 2C, magenta). This phenotype was suppressed by the ectopic SUN2-mCerlean expression, indicating that the observed motility defects were caused by SUN2 depletion. We conclude that directional nuclear migration during interphase is partially impaired by the loss of *SUN2*.

**Figure 2.**
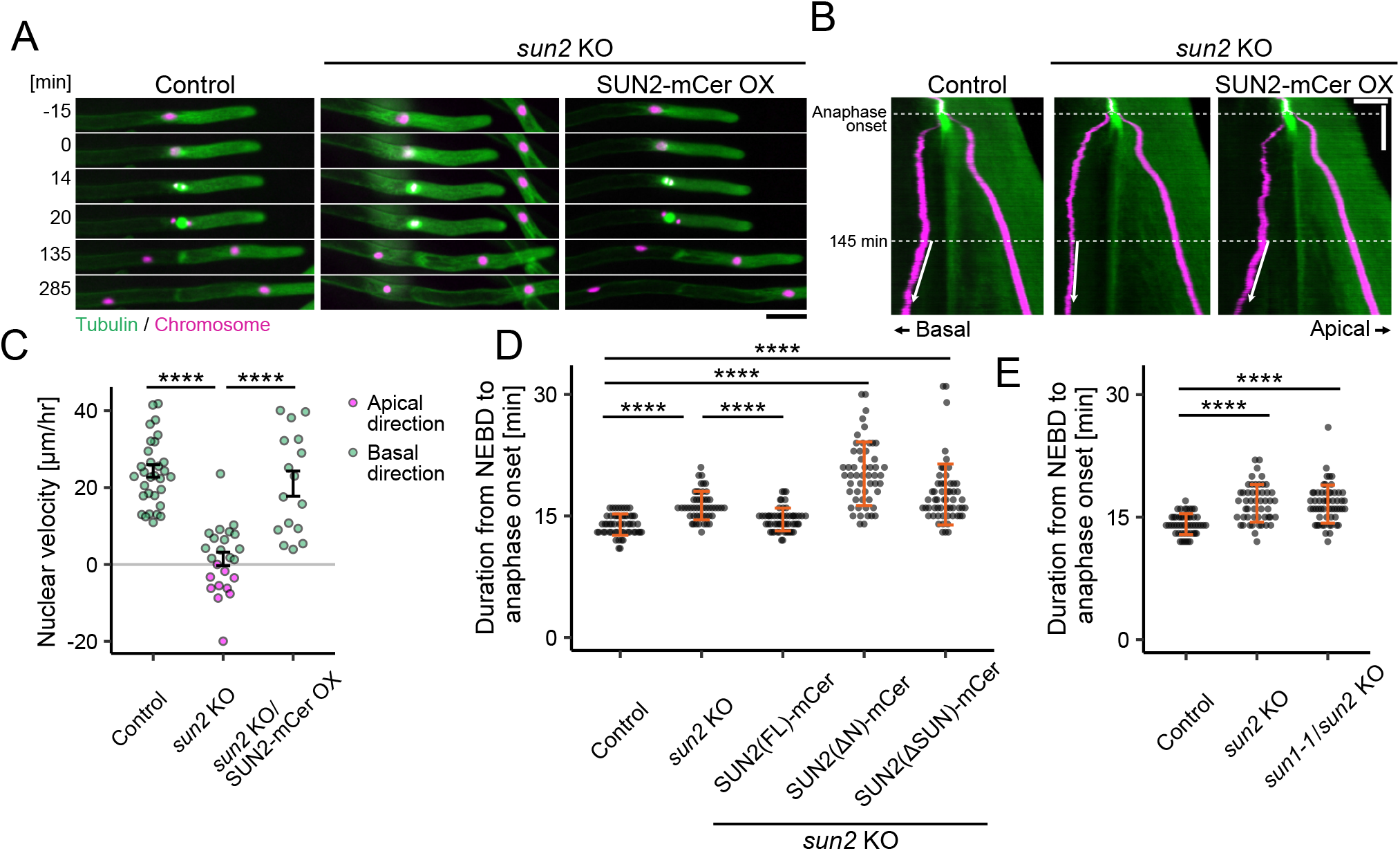
Defective nuclear migration and mitotic delay in the *sun2* KO line. (A) Nucleus dynamics in protonemal apical and subapical cells. NEBD was set to 0 min. Images were acquired with epifluorescence (wide-field) microscopy using a 10x lens. Bar, 50 µm. (B) Kymographs showing nuclear movement after cell division. The movement after 145 min is indicated by arrows. Bars, 50 µm (horizontal) and 50 min (vertical). (C) Nuclear movement rate in subapical cells 145 min after anaphase onset. The apical and basal movements were counted as negative and positive values, respectively. OX stands for overexpression. Mean ± SEM (min): 24.3 ± 1.63 (n = 32), 1.44 ± 1.76 (n = 24), 21.0 ± 3.27 (n = 16). P-values based on two-sided Tukey’s multiple comparison test: P < 0.0000001 (control vs. *sun2* KO) and P = 0.0000002 (*sun2* KO vs. *sun2* KO/SUN2 [full-length]-mCerulean). (D) Mitotic duration. Mean ± SEM (from left to right): 13.9 ± 0.178 min (n = 55), 16.3 ± 0.249 min (n = 50), 14.5 ± 0.190 min (n = 55), 20.2 ± 0.540 min (n = 53), and 17.6 ± 0.498 min (n = 57). P-values based on two-sided Steel-Dwass test: P < 0.00001 (control vs. *sun2* KO), P = 0.0001 (*sun2* KO vs. *sun2* KO/SUN2 [full-length]-mCerulean), P < 0.00001 (control vs. *sun2* KO/ SUN2DN-mCerulean), and P < 0.00001 (control vs. *sun2* KO/ SUN2DSUN-mCerulean). (E) Mitotic duration of *sun1-1*/*sun2* KO. Mean ± SEM (from left to right): 14.1 ± 0.181 min (n = 50), 16.7 ± 0.322 min (n = 51), 16.6 ± 0.308 min (n = 57). P-values based on two-sided Steel-Dwass test: P = 0.9543 (*sun2* KO vs. *sun1-1*/*sun2* KO).

### Mitosis is delayed in the absence of SUN2

In addition to nuclear motility, we identified mitotic defect in the *sun2* KO line. *P. patens* protonemal apical cells undergo highly precise mitotic cell division under laboratory culture conditions (Nakaoka et al., 2012). We confirmed this by observing 55 mitotic events in the control line: the duration from nuclear envelope breakdown (NEBD) to anaphase onset was 13.9 ± 1.32 min (± SD). This duration was significantly increased in the *sun2* KO line (16.3 ± 1.76 min; n = 50) (Fig. 2D). This phenotype was suppressed when SUN2-mCerlean was ectopically expressed (14.6 ± 1.41 min, n = 55).

To test whether the N-terminal (putative chromatin-binding) or C-terminal (putative KASH/WIP-binding) domain of SUN2 is responsible for mitotic progression, two truncated constructs were constructed and individually transformed into the *sun2* KO line. Neither construct restored mitotic duration; rather, they further extended it (Fig. 2D). These results indicated that both termini are required for rapid mitotic progression.

For unknown reasons, we were unable to obtain a moss line with complete deletion of the *SUN1* gene. Therefore, we generated *SUN1* loss-of-function mutants using CRISPR/Cas9 with different guide-RNA sets. We obtained three alleles from the background of *sun2* KO lines. In one allele, a 298 bp deletion was detected in exon2 and exon3 of *SUN1* (*sun1-1*/*sun2* KO allele) (Fig. S1C). This is likely a strong loss-of-function allele of Cter-SUN. However, the mitotic duration was not further extended compared to single *sun2* KO (Fig. 2E). These results indicate that SUN2 plays a major role in controlling cell division in protonemata.

### SUN2 is required for proper positioning of mitotic MTOC in protonemal cells

To examine the nuclear morphology and spindle/chromosome dynamics, we used high-resolution live microscopy. In the control line, the nucleus in the subapical cells was ellipsoidal during interphase, whereas it was rounder in the apical cells (Fig. 3A, B). Nuclear morphology dynamically changes during prophase in apical cells. The nucleus became ellipsoidal or diamond-shaped 10–90 min before NEBD, concomitant with the observation of the surrounding microtubule bundles (Fig. 3C, D). The microtubules applied force to the nucleus: no change in shape was observed when the microtubules were depolymerised with oryzalin (see Fig. 6B). Nuclear shape was markedly different in the *sun2* KO line. During interphase in subapical cells and prophase in apical cells, the nucleus remained round-shaped in the *sun2* KO line (Fig. 3A–D). Microtubule bundles around the NE were less prominent in the KO background during prophase and interphase (Fig. 3A, C, E). The nuclear phenotype was rescued by ectopic expression of SUN2-mCerulean. Thus, SUN2 is required for microtubule-dependent nuclear morphogenesis, which is consistent with previous observations in Arabidopsis (Oda and Fukuda, 2011; Zhou et al., 2012).

**Figure 3.**
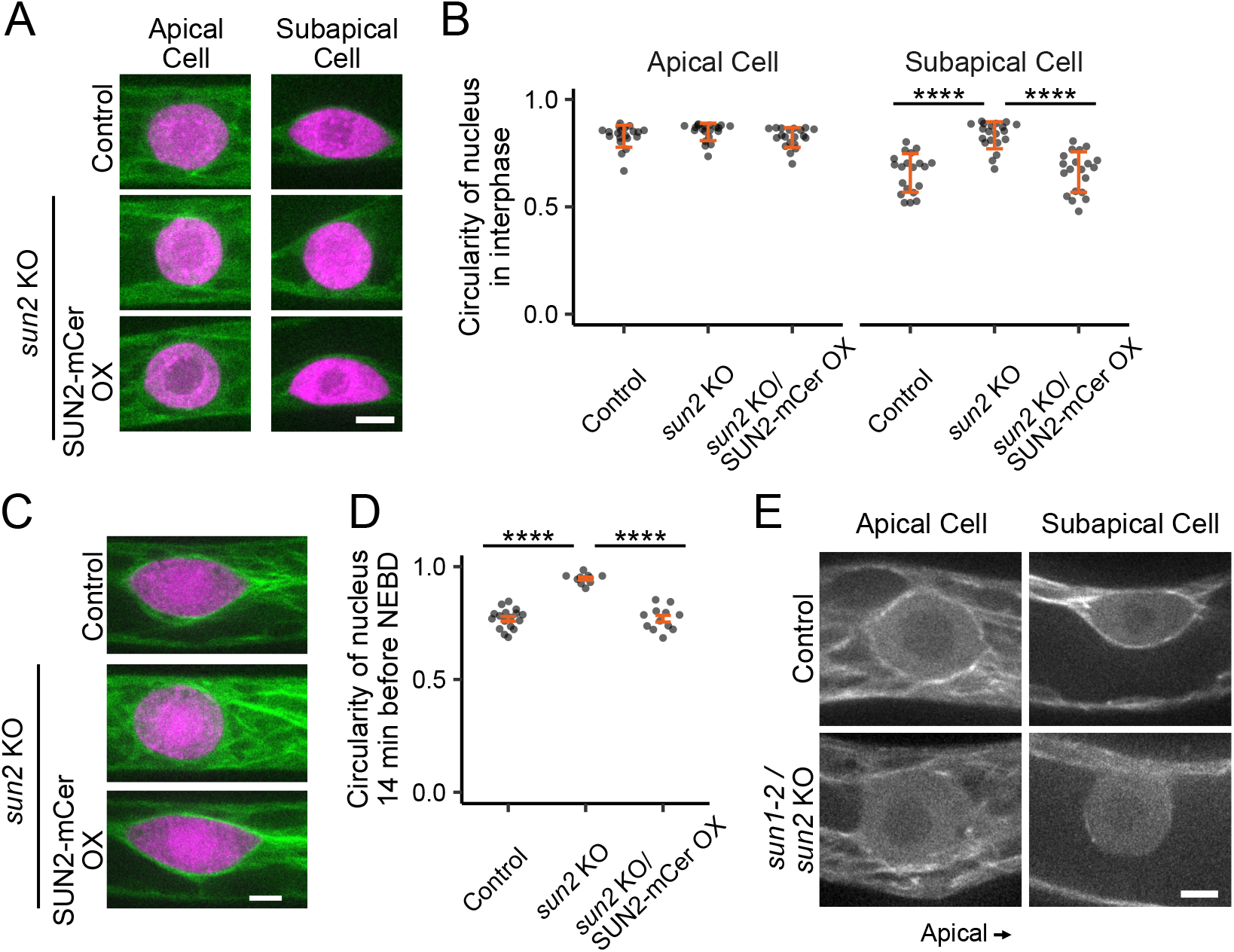
Nucleus deformation in the *sun2* KO line. (A) Shape of the nucleus in interphase. Green; microtubules. Magenta; chromosomes. (B) Circularity of the nucleus in interphase. Mean ± SEM (from left to right): Apical cells, 0.829 ± 0.0115 (n = 20), 0.849 ± 0.00904 (n = 20), 0.823 ± 0.0103 (n = 20), Subapical cells, 0.658 ± 0.0201 (n = 20), 0.833 ± 0.0141 (n = 20), 0.662 ± 0.0211 (n = 20). P-values based on two-sided Steel-Dwass test; P < 0.00001 (subapical cells: control vs. *sun2* KO), P < 0.00001 (subapical cells: *sun2* KO vs. *sun2* KO/SUN2 [full-length]-mCerulean). (C) Shape of the prophase nucleus 14 min before NEBD. Green; microtubules. Magenta; chromosomes. (D) Circularity of the nucleus 14 min before NEBD. Mean ± SEM (from left to right): 0.769 ± 0.0107 (n = 17), 0.947 ± 0.00711 (n = 10), 0.769 ± 0.0148 (n = 12). P-values based on two-sided Tukey’s multiple comparison test; P < 0.0000001 (subapical cells: control vs. *sun2* KO), P < 0.0000001 (subapical cells: *sun2* KO vs. *sun2* KO/SUN2 [full-length]-mCerulean). (E) Microtubules around the nucleus in protonemal cells in interphase Control; mCherry-α-tubulin line. Bars, 5 µm.

Just before NEBD (<10 min), the nucleus in the control apical cells transformed again to round shape, accompanied by the emergence of microtubule ‘apical cap’, which refers to the accumulation of microtubules at the apical side of the nuclear surface (Fig. 4A, Movie 2). These microtubules, as MTOC, are thought to offer a force to change nuclear morphology, move the late prophase nucleus, and serve as the initial source of spindle microtubules after NEBD (Nakaoka et al., 2012). The apical cap was infrequently observed in *sun2* KO line (Fig. 4A, Movie 2). Instead, MTOCs were detected at different positions in 21 of the 25 cells (Fig. 4B). This characteristic phenotype was suppressed by SUN2-mCerlean expression, indicating that the apical cap abnormality was caused by SUN2 protein depletion. Microtubule bundles around the NE on the basal side also reduced (Fig. 4A, Movie 2). We conclude that SUN2 is required for microtubule association with NE in late prophase.

**Figure 4.**
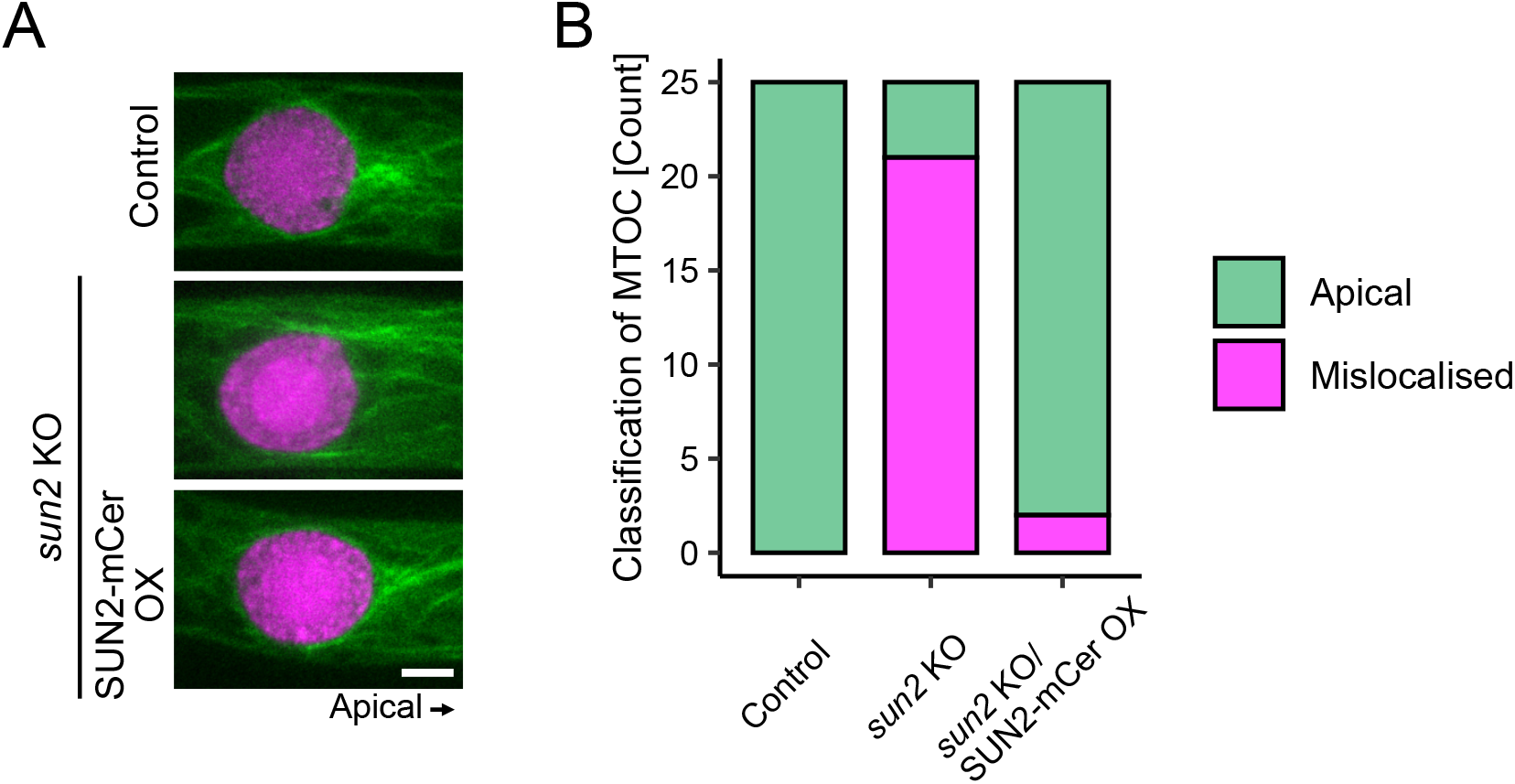
Abnormal MTOC position in prophase in the *sun2* KO line. (A) Apical cap MTOC right before NEBD. Green; microtubules. Magenta; chromosomes. Bar, 5 µm. (B) Frequency of apical NE association of MTOCs (n = 25 each).

### SUN2 is required for efficient chromosome-microtubule interaction

In addition to MTOC position, high-resolution imaging revealed differences in chromosomal dynamics during spindle assembly (Fig. 5A, Movie 2). In control cells, the nucleus rapidly migrated apically in late prophase, followed by NEBD (Fig. 5B, C [kymographs]). The apical motility of the chromosomes persisted for a few minutes after NEBD. Chromosomes on the basal side of the nucleus travelled more rapidly and over longer distances than those on the apical side, leading to chromosome congression at the spindle equator within ∼10 min (Fig. 5C). In contrast, rapid apical movement of the nucleus in late prophase and prometaphase was largely suppressed in the *sun2* KO line. Furthermore, the histone signal in the kymograph tended to extend basally after NEBD; consequently, the initiation of apical migration was delayed (red arrowheads in Fig. 5B and C). In two cases, we observed a clearly misaligned chromosome detached from the spindle body (Fig. 5A, arrowheads). However, misaligned chromosomes were eventually captured by spindle microtubules during prolonged prometaphase, which was different from kinetochore deficiency, in which chromosome congression is never achieved (Kozgunova et al., 2019). We conclude that SUN2 facilitates chromosome alignment in the spindle.

**Figure 5.**
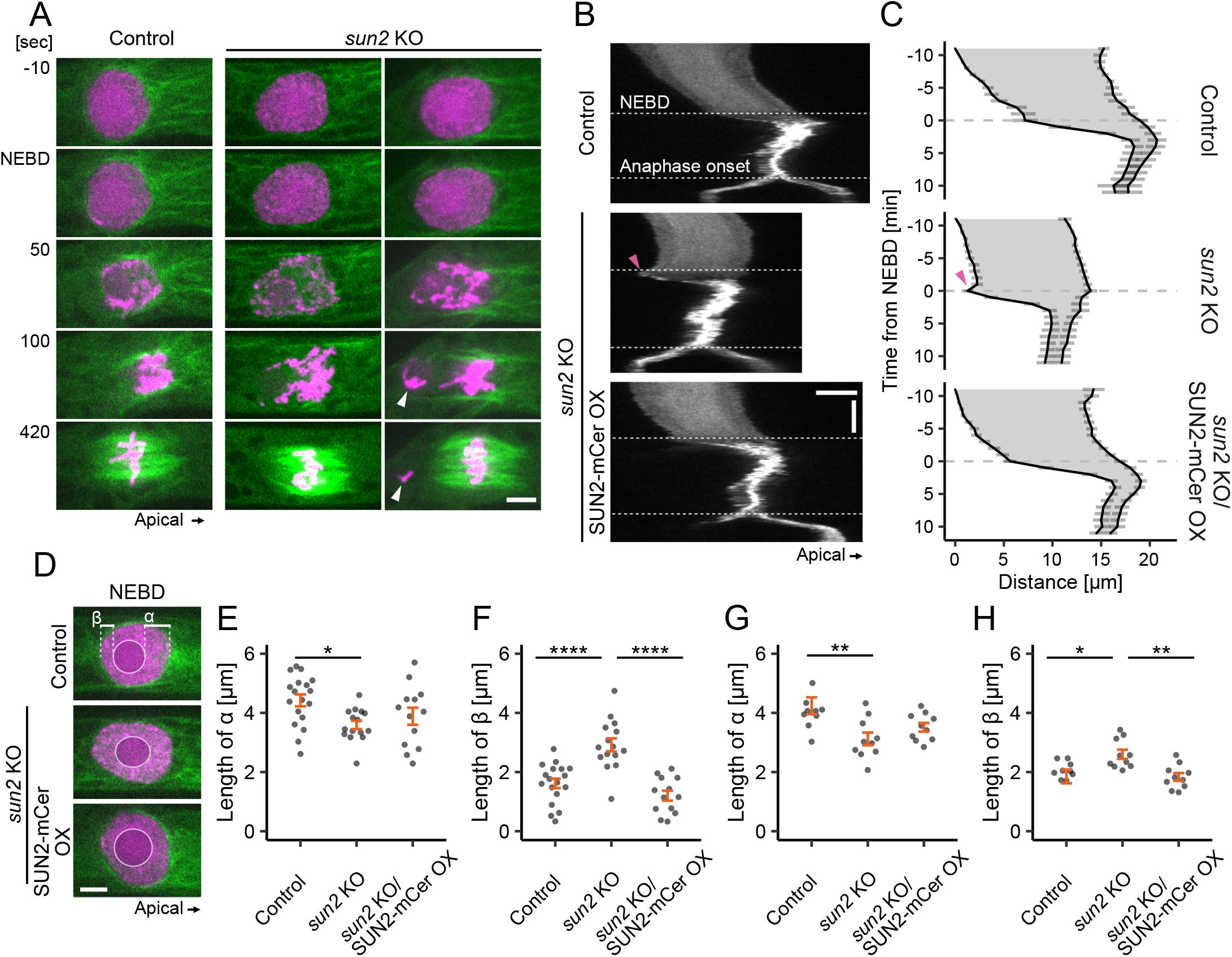
Delayed chromosome congression in the *sun2* KO line. (A) Spindle assembly and chromosome congression. Two examples are shown for *sun2* KO. The arrowhead indicates a misaligned chromosome. NEBD was set to 0 sec. Green; microtubules. Magenta; chromosomes. Bar, 5 µm. (B) Kymographs showing the dynamics of chromosome mass. Arrowhead indicates characteristic basal motility upon NEBD observed in *sun2* KO. Bar, 5 µm (horizontal) and 5 min (vertical). (C) Quantification of nuclear and chromosomal dynamics (mean ± SEM). The contour of Histone-mCherry of the kymograph signal was averaged. From top to bottom, n = 11, 7, 10. Arrowhead represents characteristic basal motility upon NEBD in *sun2* KO. (D) Position of the nucleolus (circled) at the NEBD. The distance between the nucleolus and apical edge of the nucleus (α) or basal edge of the nucleus () was quantified in (E) and (F), respectively. Bar, 5 µm. (E) Quantification of the length α in (D) at NEBD. Mean ± SEM (from left to right): 4.42 ± 0.201 (n = 18), 3.59 ± 0.142 (n = 15), 3.89 ± 0.289 (n = 13). P-values based on two-sided Tukey’s multiple comparison test; P = 0.0175847 (subapical cells: control vs. *sun2* KO), P = 0.6239412 (subapical cells: *sun2* KO vs. *sun2* KO/SUN2 [full-length]-mCerulean). (F) Quantification of the length in (D) at NEBD. Mean ± SEM (from left to right): 1.62 ± 0.159 (n = 18), 2.92 ± 0.220 (n = 15), 1.20 ± 0.168 (n = 13). P-values based on two-sided Tukey’s multiple comparison test; P = 0.0000163 (subapical cells: control vs. *sun2* KO), P = 0.0000004 (subapical cells: *sun2* KO vs. *sun2* KO/SUN2 [full-length]-mCerulean). (G) Quantification of the length α in (D) right before NEBD. Mean ± SEM (from left to right): 4.24 ± 0.284 (n = 10), 3.12 ± 0.216 (n = 10), 3.52 ± 0.146 (n = 10). P-values based on two-sided Tukey’s multiple comparison test; P = 0.0038877 (subapical cells: control vs. *sun2* KO), P = 0.4316128 (subapical cells: *sun2* KO vs. *sun2* KO/SUN2 [full-length]-mCerulean). (H) Quantification of the length in (D) right before NEBD. Mean ± SEM (from left to right): 1.85 ± 0.231 (n = 10), 2.60 ± 0.156 (n = 10), 1.84 ± 0.129 (n = 10). P-values based on two-sided Steel-Dwass test; P = 0.0116 (subapical cells: control vs. *sun2* KO), P = 0.0076 (subapical cells: *sun2* KO vs. *sun2* KO/SUN2 [full-length]-mCerulean).

During mitosis observation, we noticed that chromosomal organisation within the nucleus was skewed in late prophase in the *sun2* KO line. The fluorescent histone marker visualised the nucleosome-based chromosomes and RNA-rich nucleolus as distinct signals (Fig. 5D). The nucleolus is usually located at the centre of the nucleus during interphase and early prophase. However, just before or after NEBD, chromosome masses were more enriched on the apical side, and consequently, the nucleolus was positioned closer to the basal edge of the nucleus in control cells (Fig. 5D–H). In contrast, the nucleolus remained medially localised and a clear bias in chromosome distribution was not observed in the *sun2* KO line. These observations suggest that SUN2 mediates trans-NE microtubule-chromosome interactions at the onset of NEBD.

### Asymmetric SUN2 distribution during apical MTOC assembly

As expected, endogenous SUN2 tagged with mNeonGreen (mNG) was uniformly localised to the NE during interphase (Fig. 6A). In contrast, time-lapse microscopy and signal quantification indicated that SUN2 localisation was asymmetric in late prophase; SUN2-mNG was more enriched on the apical side, partially overlapping the microtubule apical cap (Fig. 6B, C).

To test whether the asymmetric distribution was dependent on microtubules, we depolymerised the cytoplasmic microtubules with oryzalin, followed by time-lapse microscopy. Quantification of signal intensity showed no apical accumulation of SUN2 in late prophase under these conditions (Fig. 6B, C). In contrast, the depolymerisation of actin filaments with latrunculin A did not disrupt the asymmetric distribution of SUN2 (Fig. 6B, C). These results indicate that apical enrichment of SUN2 and microtubules is mutually dependent.

During spindle assembly, SUN2-mNG initially showed punctate signals on the spindle (Fig. 6D). The number of signals gradually decreased, and the spindle at metaphase was cleared.

**Figure 6.**
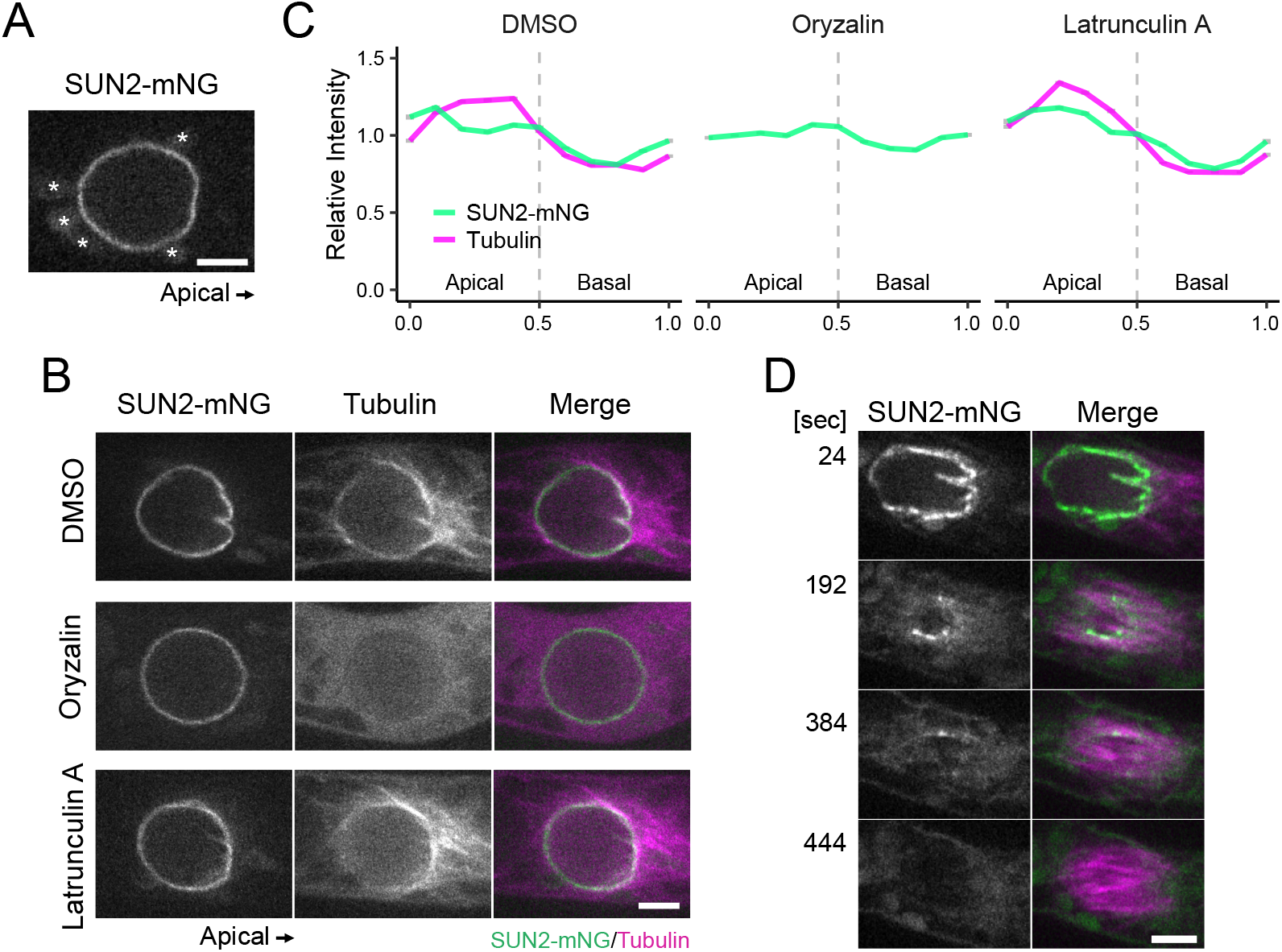
Localisation of SUN2 in early mitosis. (A) SUN2-mNeonGreen (mNG) localisation in interphase. Asterisks indicate autofluorescent chloroplasts. (B) SUN2 localisation in late prophase with or without cytoskeleton drugs. (C) Quantification of the SUN2-mNG distribution along the NE. Apical enrichment was disrupted by oryzalin treatment but not by Latrunculin A. From left to right, n = 16, 14, 15 cells. (D) SUN2 localisation during spindle assembly. Time 0; NEBD. Bars, 5 µm.

### SUN2 controls MTOC position and microtubule-NE interaction in the gametophore initial

The gametophore in Physcomitrella is the leafy shoot, which develops from protonemal filaments. In the first stage of gametophore development, stem cells undergo a type of asymmetric division distinct from that of the protonemata (Harrison et al., 2009; Kofuji and Hasebe, 2014). We examined the role of Cter-SUN in this cell type. Similar to the protonemata, a portion of the microtubules surrounded the NE during prophase in the gametophore initial cells (Fig. 7A, red arrowhead). We observed that the signals of NE-surrounding microtubules decreased in the absence of SUN2 (Fig. 7A). In addition, in this system, the microtubule cloud, or also called regional MTOC ‘gametosome’, emerges at the apical cytoplasm and functions as the dominant microtubule nucleation site (Kosetsu et al., 2017) (Fig. 7A, green arrowhead). In control cells, gametosomes appeared in the apical cytoplasm in prophase, and the line connecting the nuclear and gametosome centres was nearly parallel to the long axis of the cell (Fig. 7B, C). In the *sun2* KO line, gametosomes were formed normally. However, its position relative to the nucleus was more variable (Fig. 7C). Thus, SUN2 plays a similar role in the gametophore initial cell in terms of microtubule-NE association and cytoplasmic MTOC (i.e. gametosome) positioning. The gametosome dictates spindle orientation, and consequently, the division plane in gametophore initial cells (Kosetsu et al., 2017).

**Figure 7.**
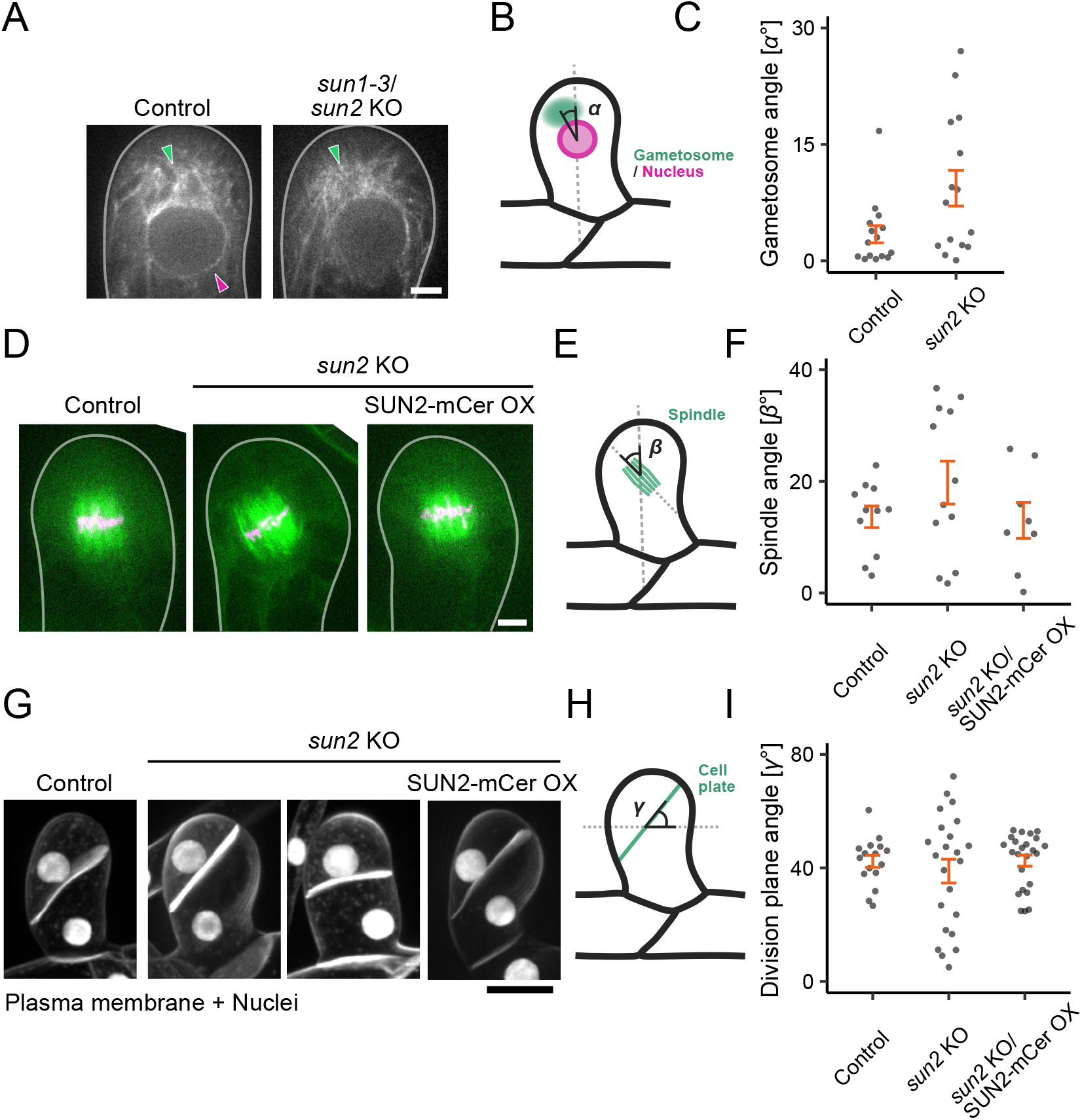
Division plane misorientation in gametophore initial cells of the *sun2* KO line. (A) Microtubule clouds called ‘gametosome’ (green arrowheads) appeared in the apical cytoplasm in prophase in the gametophore initial cell. The magenta arrowhead indicates accumulation of microtubules around the basal NE. Control; mCherry-α-tubulin line. Bar, 5 µm. (B) Scheme of the angle (*a*) measurements. Green; gametosome. Magenta; nucleus. (C) Quantification of the relative positions of gametosomes in cells. The angle *a* defined in (B) was measured. Mean ± SEM (from left to right): 3.43 ± 1.11 (n = 15), 9.35 ± 2.29 (n = 15). P-values based on two-sided Mann-Whitney U test: P = 0.05022 (control vs. *sun1-3*/*sun2* KO). (D) Metaphase spindles in the gametophore initial cell. Green; microtubules. Magenta; chromosomes. Bar, 5 µm. (E) Spindle angle (*{3*) measurement scheme. (F) Quantification of metaphase spindle orientation The angle *{3* defined in (E) was measured. Mean ± SEM (from left to right): 13.6 ± 1.93 (n = 11), 19.7 ± 3.84 (n = 12), 13.0 ± 3.21 (n = 8). P-values based on two-sided Steel-Dwass test: P = 0.8454 (control vs. *sun2* KO), P = 0.6253 (subapical cells: *sun2* KO vs. *sun2* KO/SUN2 [full-length]-mCerulean). (G) The plasma membrane was visualised using FM4-64 staining in the gametophore initial cell after cell division. Bar, 20 µm. Scheme of division plane angle (*y*) measurement. (H) Quantification of division plane angle in cells. The angle *y* defined in (H) was measured. Mean ± SEM (from left to right): 42.3 ± 2.14 (n = 16), 38.9 ± 4.19 (n = 23), 42.6 ± 1.96 (n = 24). (I) P-values based on two-sided Steel-Dwass test: P = 0.9995 (control vs. *sun2* KO), P = 0.8988 (subapical cells: *sun2* KO vs. *sun2* KO/SUN2 [full-length]-mCerulean).

Consistent with the variable positions of the gametosome, the orientation of the metaphase spindle (Fig. 7D–F) and cell plate (Fig. 7G–I) were also variable in the *sun2* KO line. These results suggest that SUN2-mediated linkage between NEs and microtubules is required for proper orientation of the mitotic spindle and, thereby, that of the division plane.

## Discussion

The contribution of nuclear membrane proteins to cellular events in plants remains poorly understood. In this study, we found that the SUN2 protein in *P. patens* not only plays a well-established role in nuclear shaping and positioning but also facilitates chromosome alignment during mitosis. A series of live imaging supports a model in which SUN2 mediates the interaction between MTOC and the nucleus during mitotic prophase, enabling the efficient association of microtubules with chromosomes during spindle assembly.

### Physcomitrella SUN2 couples NE with microtubules

The loss of centrosomes is a striking evolutionary event in plant lineages (Buschmann and Zachgo, 2016). Several types of acentrosomal MTOCs have been developed as centrosome substitutes (Buschmann et al., 2016; Lloyd and Chan, 2006; Naramoto et al., 2022; Yi and Goshima, 2018). In some cases, the proteins required for MTOC formation have been identified (Liu and Lee, 2022). However, little is known about the spatial control of acentrosomal MTOCs in plants. The apical cap of the moss protonema represents a form of MTOC that requires γ-tubulin and TPX2 for formation and functions in early phase of mitosis as the major source of spindle microtubules (Kozgunova et al., 2022; Nakaoka et al., 2012). The current study demonstrates that SUN2 is required for the association between MTOCs and NE. A plausible mechanism is that SUN2 links cytoplasmic MTOC to the NE through unidentified KASH/WIP/WIT proteins. Microtubule-dependent apical enrichment of SUN2 during prophase is consistent with this notion. The function of SUN2 may not be limited to the prophase; the interphase nucleus was deformed concomitant with a reduction in the surrounding microtubules in the *sun2* KO line. In animals, the SUN protein forms a central part of the LINC complex, which connects the cytoskeleton, including microtubules and actin, with nuclear laminae and chromosomes across the NE (Gundersen and Worman, 2013; Mejat and Misteli, 2010). In Arabidopsis, SUN mediates actin-NE interactions via WIP/WIT and myosin XI-i (Tamura et al., 2013). Our results indicated that the microtubule-linking function of SUN is preserved in plants.

### SUN2 facilitates chromosome-microtubule interaction during mitosis

The skewed distribution of prophase chromosomes in the *sun2* KO line, exemplified by the mispositioned nucleolus, suggests that SUN2 also mediates the coupling of chromosomes with microtubules. It is conceivable that the proximity of microtubule bundles to chromosomes facilitates their interactions with NEBD. Therefore, we propose that microtubule mispositioning outside the nucleus and chromosome disorganisation within the prophase nucleus additively delay microtubule-chromosome interactions during spindle assembly. However, it is not ruled out that SUN2 also actively participates in the spindle assembly process during early prometaphase, for example, by removing nuclear membrane remnants in the spindle matrix (Turgay et al., 2014).

Our model is reminiscent of what is known about *S. pombe* Sad1, the founder of the SUN family (Hagan and Yanagida, 1995). In yeast, NE does not completely disassemble during mitosis. Instead, the insertion of SPB, i.e., point MTOC, into NE is critical for spindle assembly during mitosis (Fernandez-Alvarez et al., 2016; Jaspersen et al., 2006; Mejat and Misteli, 2010). Sad1 is localised to the SPB throughout the cell cycle and links to the centromeres of each chromosome during interphase. Mutant analysis has shown that pre-mitotic contact between centromeres and SPB is required for proper initiation of mitosis (Fernandez-Alvarez et al., 2016). Thus, the SUN-mediated trans-NE connection between MTOC and chromosomes may be a conserved mechanism that guarantees robust cell division, despite a significant morphological difference between the SPB (point MTOC) and apical cap (regional MTOC).

### Limitations of the study

Compared with the known loss-of-function phenotype of SUN in animal cells, the phenotypes observed in this study were mild. For example, we did not detect defects in nuclear migration in the apical cells of *sun2* KO or *sun1-1 / sun2* KO. Similarly, in Arabidopsis, *sun1* KO / *sun2* knockdown line showed no dramatic developmental or fertility defects under laboratory conditions (Oda and Fukuda, 2011; Zhou et al., 2012). We speculate that this is due to the presence of the intact mid-SUN in the mutants, which may also act as a linker between microtubules and NE (Graumann et al., 2014; Meier et al., 2017). Despite several attempts using different constructs, we could not obtain KO or mutant alleles for *SUN3*; SUN3 might be required for essential processes in cell proliferation. Similarly, the Arabidopsis mid-SUN triple mutant is lethal (Graumann et al., 2014). A comprehensive loss-of-function analysis of Cter- and mid-SUN would be an interesting topic for future research.

The types of plant MTOCs that require SUN for localisation are another outstanding question. In seed plants, many cell types develop a specialised regional MTOC called ‘polar cap’ or ‘pro-spindle’ in late prophase, which caps both apical and basal sides of the NE (Liu and Lee, 2022; Smirnova and Bajer, 1998). AtSUN1 and AtSUN2 are abundantly localised at the polar cap (Oda and Fukuda, 2011; Tatout et al., 2014). It would be interesting to revisit the *sun* mutants and examine whether SUN mediates the association between NE and this type of MTOC.

## Materials and methods

The majority of the methods used in this study were identical to those described in our recent studies (Ta et al., 2023; Yoshida et al., 2023).

### *P. patens* culture and transformation

All strains in this study were derived from the Gransden ecotype of *Physcomitrium* (*Physcomitrella*) *patens* (Ashton and Cove, 1977). *P. patens* culture and transformation protocols followed were as described by (Yamada et al., 2016). Briefly, mosses were regularly cultured on BCDAT plates at 25 °C under continuous light illumination. A standard polyethylene glycol (PEG)-mediated method was exploited for transformation.

Prior to transformation, sonicated protonemata were cultured on BCDAT agar medium for 5–6 d. Transgenic lines were selected using corresponding antibiotics. Line confirmation was conducted through visual inspection followed by genotyping PCR (Fig. S1, Table S4). Sequencing was performed to confirm the CRISPR mutant lines. The lines generated in this study are listed in Table S1.

### Plasmid construction

The plasmids and primers used in this study are listed in Tables S2 and S3, respectively. CRISPR targets with high specificity were manually selected in the first three exons of *SUN1* gene, and the regions near the start or stop codons of *SUN2* gene. All target sequences were synthesised and ligated into the *Bsa*I site of pPY156, which is based on pCasGuide/pUC18 and contains a hygromycin-resistant cassette (Lopez-Obando et al., 2016; Yi and Goshima, 2020). For endogenous tagging via homologous recombination, the plasmid was constructed using the In-Fusion HD Cloning Kit (Takara); 1–2 kb sequences of the 5’ and 3’ ends of the genes of interest flanked the fragment that consisted of an in-frame linker, mNeonGreen (mNG) tagged with FLAG, or mCherry coding sequence, and G418 or blasticidin S resistant cassette. The mNG codon was optimised for expression in Arabidopsis. For all the rescue experiment, the *SUN2* coding sequence was amplified from the moss cDNA library (full-length, truncation, mutant) and ligated into the pENTR/D-TOPO vector containing the in-frame linker, Cerulean-coding sequence, followed by the Gateway LR reaction (Invitrogen) into a vector containing the *P. patens EF1α* promoter, nourseothricin resistance cassette, and 1-kb sequences homologous to the *PTA1* locus.

### Moss growth assay

The 5–7-day-old sonicated protonemata with similar sizes were inoculated to the BCDAT plate. Two plates each containing 25 pieces of inoculated protonemata were made for each strain. After 20 d of incubation under the continuous light, images of overall moss or gametophores were acquired using a C-765 Ultra Zoom digital camera (Olympus) or SMZ800N, respectively.

### Microscopy

Time-lapse microscopy was performed as described by (Nakaoka et al., 2012). Briefly, in the long-term time-lapse imaging experiments for the observation of protonemal cells, the protonemata were cultured on thin layers of BCD agarose in 6-well glass-bottom dishes for 5–7 d. Wide-field, epifluorescence images were acquired with a Nikon Ti microscope (10× 0.45 NA lens, Zyla 4.2P CMOS camera (Andor), Nikon Intensilight Epi-fluorescence Illuminator) at intervals of 1 min (no z-stacks). For high-resolution imaging, protonemata were inoculated onto the agar pad in a 35 mm glass-bottom dish, followed by culturing for 5–7 d. Confocal imaging was performed with a Nikon Ti microscope attached to a CSU-X1 spinning-disc confocal scanner unit (Yokogawa), EMCCD camera (ImagEM, Hamamatsu), and three laser lines (561, and 488 nm). 100× 1.45 NA lens was used for most experiments related to protonemal and gametophore cell divisions. For the quantification of gametophore division plane angle, 40× 1.30 NA lens was used. To induce the gametophore cells, protonemal cells were treated with 2-isopentenyladenine (2iP) for 5–10 min, 20–22 h before imaging (Kosetsu et al., 2017). Stock solution of oryzalin, latrunculin A, FM4-64 and 2iP in DMSO was diluted with distilled water to working concentrations of 10 µM oryzalin, 25 µM latrunculin A, 10 µM FM4-64 and 1 µM 2iP. Prior to drug addition, the protonemal tissue on the agarose pad was preincubated in water for 1 h for absorption. After water removal, 0.3 mL of the drug solution was added, and image acquisition was started 5–10 min later. DMSO was used as a control. Imaging was performed at 22–25 °C in the dark.

### Image data analysis

All raw data processing and measurements were performed using the Fiji software.

#### Moss growth on the culture plate

The images of the moss on the culture plate were outlined automatically, and the area was measured using Fiji.

#### Mitotic duration

For caulonemal apical cells, time-lapse images were obtained every 1 min using an epifluorescence (wide-field) microscope and a 10× 0.45 NA lens, and the duration between NEBD and the anaphase onset was measured.

#### Nuclear velocity

For caulonemal apical cells, time-lapse images were obtained every 1 min using an epifluorescence (wide-field) microscope and a 10× 0.45 NA lens. A kymograph of chromosomes was generated along the dividing cell. To obtain nuclear velocity, the inclination of the nuclear signal in the subapical cell 145 min after anaphase onset was manually measured with Fiji.

#### Nuclear circularity

For caulonemal apical and subapical cells in interphase, the images of the nuclei were obtained with z-stacks at 1 µm intervals for a range of 9 µm using a spinning-disc confocal microscope and a 100× 1.45 NA lens. The z-stack images were processed by maximum z-projection. The nuclei were outlined automatically, and the circularity of the nucleus was measured with Fiji. For caulonemal apical cells in mitotic prophase, time-lapse images were obtained every 2 min with z-stacks at 1 µm intervals for a range of 7 µm using a spinning-disc confocal microscope and a 100× 1.45 NA lens. The circularity of the nucleus at the best focal plane was measured manually with Fiji.

#### Chromosome dynamics during cell division

For caulonemal apical cells, time-lapse images were obtained every 10 s with z-stacks at 2 µm intervals for a range of 4 µm using a spinning-disc confocal microscope and a 100× 1.45 NA lens. The best focal plane was selected for analysis. A kymograph of chromosome mass was generated along the spindle pole-to-pole axis. For quantification, the chromosome mass on the kymograph was manually outlined with Fiji and, for each timepoint, the distance from the basal edge of the nucleus (set at 11 min before NEBD) was calculated.

#### Distance between the nucleolus and nuclear edge

For caulonemal apical cells, time-lapse images were obtained every 10 s with z-stacks at 2 µm intervals for a range of 4 µm using a spinning-disc confocal microscope and a 100× 1.45 NA lens. The best focal plane in prophase was selected for analysis. The radius of the nucleus, the distance from the nuclear apical edge to the nucleolus centre, and the diameter of the nucleus were measured manually with Fiji.

#### Distribution of SUN2-mNG along the nuclear membrane

For caulonemal apical cells, time-lapse images were obtained every 30 s with z-stacks at 1.5 µm intervals for a range of 3 µm using a spinning-disc confocal microscope and a 100× 1.45 NA lens. The best focal plane was selected for analysis. The intensity of SUN2-mNG and the microtubules along the nucleus was measured just before NEBD, and the background intensity of each image was subtracted. The intensity of each pixel on the drawn line (3-pixel width) was divided by the mean intensity of the entire length of the line to get the relative intensity. The cells were divided into 10 sections, and the average relative intensity of each section is displayed.

#### Gametosome position

For gametophore initial cells, time-lapse images were obtained every 30 s with z-stacks at 1 µm intervals for a range of 6 µm using a spinning-disc confocal microscope and a 100× 1.45 NA lens. The best focal plane was selected for analysis. The relative angle between the line connecting the nuclear centre to gametosome centre and the long axis of the cell was calculated.

#### Spindle orientation

For gametophore initial cells, time-lapse images were obtained every 30 s with z-stacks at 3 µm intervals for a range of 12 µm using a spinning-disc confocal microscope and a 100× 1.45 NA lens. The best focal plane was selected for analysis. The angle between the long axis of the spindle and the long axis of the cell was calculated.

#### Division plane orientation

Gametophore initial cells were stained by FM4-64 prior to the imaging. The images of gametophore initial cells were obtained with z-stacks at 2.5 µm intervals for a range of 25 µm using a spinning-disc confocal microscope and a 40× 1.30 NA lens. The z-stack images were processed by maximum z-projection with Fiji, and the relative angle between the division plane and the long axis of the cell was calculated.

### Statistical analysis

The Shapiro-Wilk test was used for all samples to check for normality. If the sample was assumed to be normally distributed, the F-test (two groups) or Bartlett’s test (multiple groups) was conducted to test homoscedasticity. If the samples had a normal distribution and equal variance, Student’s *t*-test (two groups) or Tukey’s multiple comparison test (multiple groups) was used. If the samples had a normal distribution but not equal variance, Welch’s two-sample *t*-test (two groups) or the Games-Howell test (multiple groups) was used. If the samples did not have a normal distribution, Mann-Whitney U test (two groups) or Steel-Dwass test (multiple groups) was used. All statistical analyses were performed using R software. Obtained P values are denoted as follows: *, P < 0.05; **, P < 0.01; ***, P < 0.001; and ****, P < 0.0001. Data from multiple experiments were combined because of insufficient sample numbers in a single experiment unless otherwise stated.

## Accession numbers

The gene sequences used in this study are available in Phytozome under the following accession numbers: *SUN1* (Pp3c7_4170), *SUN2* (Pp3c11_22530), *SUN3* (Pp3c21_2240), *SUN4* (Pp3c18_19540).

## Acknowledgements

We are grateful to Maya Hakozaki or providing moss lines; Chiemi Koketsu and Rie Inaba for media preparation. This work was funded by the Japan Society for the Promotion of Science KAKENHI (18KK0202, 22H04717, and 22H02644 to GG). MWY is a recipient of the Japan Society for the Promotion of Science pre-doctoral fellowship. The authors declare no competing interests.

## Movie legends

**Movie 1. Defective nuclear migration and mitotic delay in the *sun2* KO line**

Time-lapse video of protonemal cells expressing GFP-tubulin (green) and histone H2B-mCherry (magenta). Movies were acquired using an epifluorescence (wide-field) microscope in a single focal plane (10× 0.45 lens). Bar, 50 µm.

**Movie 2. Abnormal MTOC position in prophase and delayed chromosome congression in the *sun2* KO line**

Time-lapse video of microtubules (GFP-tubulin, green) and chromosomes (histone H2B-mCherry, magenta) in protonemal apical cells. Movies were acquired using a spinning-disc confocal microscope. Bar, 5 µm.

## Supplemental tables

**Table S1. Moss lines used in this study**

**Table S2. Plasmids used in this study**

**Table S3. Primers used for plasmid construction and sequencing**

**Table S4. Primers used for genotyping PCR**

**Figure S1.**
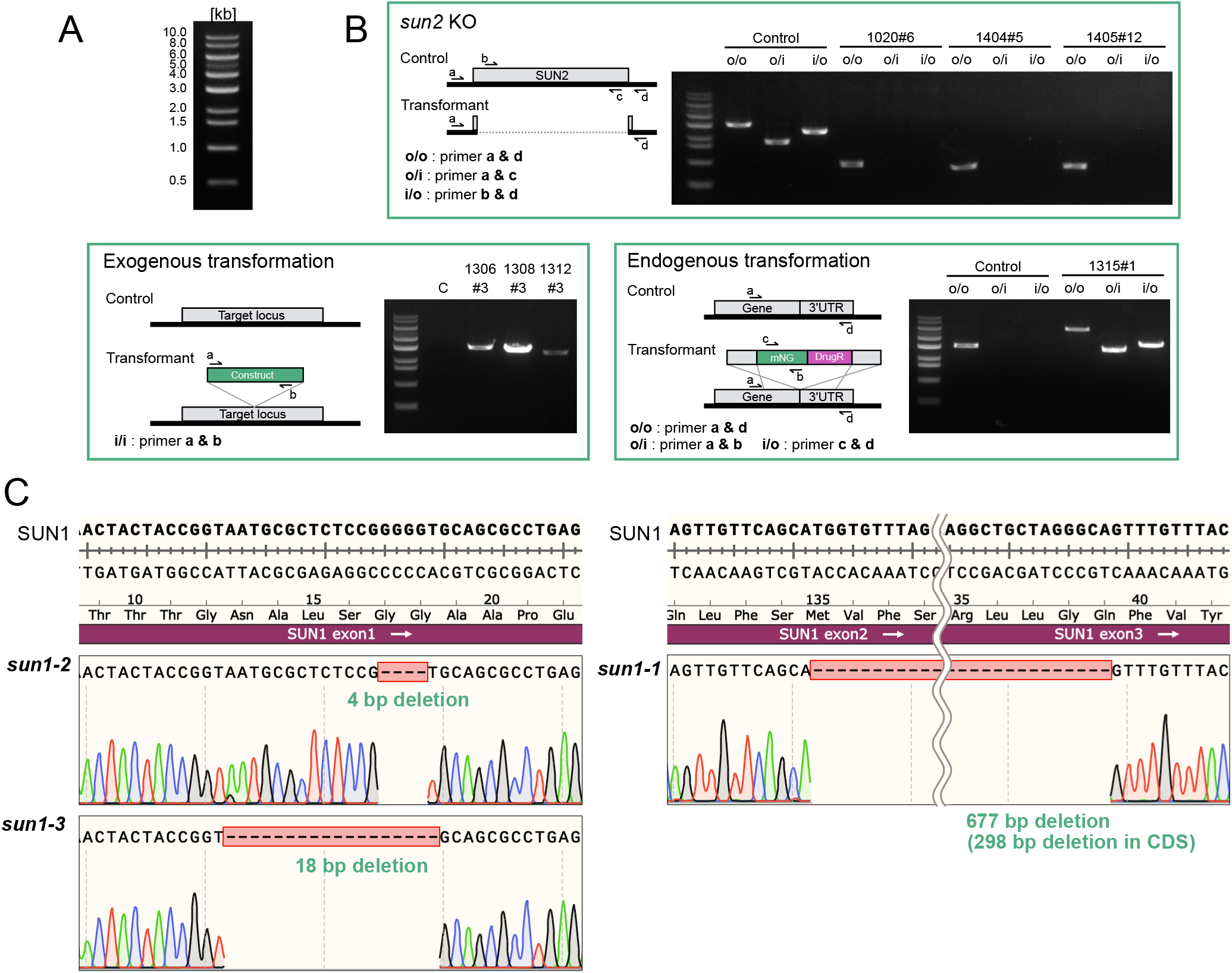
Confirmation of the moss lines established in this study. (A) Band-size marker used in DNA gel electrophoresis. (B) Genotyping PCR strategy (left) and PCR results (right) for the moss lines established in this study. Gene disruption using CRISPR/Cas9 technology (*sun2* KO), exogenous integration (rescue constructs), and C-terminal tagging (SUN2-mNG) are shown in separate boxes. The number in each lane indicates the line ID, and the genotype of each ID is listed in Table S1. “C” indicates the control (parental line). Primers used are listed in Table S4. (C) Sequencing revealed base pair deletions in the *sun1* mutants used in this study (displayed in SnapGene sequence files).

